# X-MyoNET: Biometric Identification using Deep Processing of Transient Surface Electromyography

**DOI:** 10.1101/2021.11.30.470688

**Authors:** Qin Hu, Alireza Sarmadi, Paras Gulati, Prashanth Krishnamurthy, Farshad Khorrami, S. Farokh Atashzar

## Abstract

The rapid development of the Internet and various applications such as the Internet of Medical Things (IoMT) has raised substantial concerns about personal information security. Conventional methods (e.g., passwords) and classic biological features (e.g., fingerprints) are security deficient because of potential information leakage and hacking. Biometrics that express *behavioral features* suggest a robust approach to achieving information security because of the corresponding uniqueness and complexity. In this paper, we consider identifying human subjects based on their transient neurophysiological signature captured using multichannel upper-limb surface electromyography (sEMG). An explainable artificial intelligence (XAI) approach is proposed to process the internal dynamics of temporal sEMG signals. We propose and prove the suitability of “transient sEMG” as a biomarker that can identify individuals. For this, we utilize the Gradient-weighted Class Activation Mapping (Grad-CAM) analysis to explain the network’s attention. The outcome not only decodes and makes the unique neurophysiological pattern (i.e., motor unit recruitment during the transient phase of contraction) associated with each individual visualizable but also generates an optimizing two-dimensional (2D) spectrotemporal mask used to significantly reduce the size of the model and the trainable parameters. The resulting mask selectively and systematically samples the spectrotemporal characteristics of the users’ neurophysiological responses, discarding 40% of the input space while securing the accuracy of about 74% with much shallower neural network architecture. In the systematic comparative study, we find that our proposed model outperforms several state-of-the-art algorithms. For broader impacts, we anticipate our design of a compact, practical, interpretable, and robust identification system that requires only a minimal number of gestures and sensors (only 7% of the entire data set) to be a starting point for small and portable identification hardware.

## I. Introduction

The rapid development of the Internet has contributed to the accelerated growth of several research fields, including medical technologies. An example is the Internet of medical things, which allows remote assessment and monitoring of several medical conditions. However, this has imposed risks on personal information, such as medical records [1]–[4]. The conventional methods such as Personal Identification Number and Password have been shown security deficient due to the possibility of information leakage, breaches, and counterfeits [5]–[7]. On a larger scale, there have been several reports on governmental systems and credit agencies that were compromised, exposing the information of millions of employees and customers [8]–[10]. Biological-featured methods that extract the physical characteristics of human bodies, such as features of the face, fingerprint, and iris, have been proposed as an alternative to protect information privacy [11]–[14]. However, these physiological methods are susceptible to hacking, especially since technological advances allow for duplicating three-dimensional face models using 3D printers, hacking fingerprints through latex gloves, and copying the corresponding biological features using artificial iris contact lenses [15]–[17].

As a result, there is a need for producing new means of biometrics that provide a higher level of personalization and reduce the risk of hacking. Biometrics that express *behavioral features* such as Electromyography, which is the electrical manifestation of muscle contraction [18], [19], suggest a solid approach to achieve information security robustness by the corresponding uniqueness. This is because neurophysiological responses (such as those captured by sEMG) are unique to the user and inherently complex in nature (that makes forgeries and falsifications exceedingly difficult). However, it should be noted that due to neurophysiological complexity and potential context-based variability, even for one individual, fundamental research is needed to generate techniques that can robustly and consistently detect the underlying sEMG “signature” as biomarkers. This is the focus of the current paper.

A few relatively recent works have been conducted in the literature regarding sEMG-based human identification, motivated by the unique characteristics of this biosignal to prevent personal information leakage, spoofing attacks, and identity theft. The conventional machine-learning approaches that are based on extracting temporal and spectral features from sEMG are the most commonly-used methods [20]–[23]. Examples of temporal features include Mean Absolute Value (MAV), Variance (VAR), Number of Zero-crossings (NZC), Log Detector (LD), Root Mean Square (RMS), Waveform Length (WL), Integral of the EMG (IEMG), and Difference Absolute Standard Deviation Value (DASDV). Examples of Spectral features are Median Frequency, Mean Frequency, Fast Fourier Transform (FFT) features, and Discrete Wavelet Transform (DWT) features. The extracted features are then fed into conventional classifiers such as Support Vector Machines (SVMs) and Linear Discriminant Analysis (LDA) to identify individuals.

One of the main limitations of the aforementioned studies is the simplicity of the extracted features and models. Thus, most of such efforts were conducted on small data sets, making generalized identification systems less achievable. In this regard, due to the variability, nonlinearity, and complexity of sEMG, it is imperative to test the capacity of this biosignal on a larger number of individuals and gestures to detect the unique underlying features. In recent decades, researchers have leveraged the powerful feature extraction capability of Deep Learning (DL) models to solve complex tasks. However, few of them exploited these models in human identification. In [24], the denoised sEMG signals were fed into a Convolutional Neural Network (CNN) to minimize data preprocessing and let the model learn the underlying neurophysiological patterns on its own. In addition to the above-mentioned studies, some researchers have recently converted raw sEMG signals into 2D spectrograms, concurrently analyzing temporal and spectral muscle behaviors and potentially extracting high-dimensional information for generating biomarkers. This concept has been investigated in [15], where Continuous Wavelet Transform was used in conjunction with a CNN architecture.

Even though the recent use of deep neural networks may suggest good performance, the existing works suffer from low diversification regarding subjects and hand gestures, raising concerns about generalization to a higher number of people and different gestures. Also, the existing recent DL research in personal identification cannot explain the attention of the neural network, raising concerns about the black-box modeling, including the biases in the data set, which can be challenging for identification tasks and can raise concerns about system attacks. Moreover, the use of large spectrotemporal input spaces results in computationally inefficient models, challenging practicality in terms of the size of the training set and the implementation of small and portable identification hardware.

Motivated by the points mentioned above, in this paper, we propose an identification method that uses an explainable CNN-based framework with the optimized input size derived based on the Grad-CAM attention-based analysis of the system. Our identification method is compact, practical, interpretable, and robust. It only requires a training set of less than 10% gesture and sensor data from the Ninapro Database 2 (DB2) and achieves a good accuracy of over 80%. The proposed optimal number of gestures and sensors derived from Gaussian Mixture Model (GMM) are sufficient to capture the underlying neurophysiological features associated with each subject. As mentioned earlier, DL models have been usually deemed a black box because they only show the final predicting results but not the evidence (on the inputs) for making predictions. In this paper, for the first time, we exploit explainable artificial intelligence (XAI) to interpret the proposed CNN model’s attention and visualize each subject’s extracted underlying neurophysiological patterns using Grad-CAM in sEMG-based personal identification. Besides understanding segments of spectrotemporal grids that encapsulate most of the information for identification, this explainability analysis also helps generating a 2D spectrotemporal segmenting mask. The application of the proposed mask cuts out trivial parts of the input grid, which has the minimum information for identification, further shrinking the input space and reducing the model complexity. The *<u>six main contributions</u>* of this paper are as follows:

### Contribution 1

This is the first paper that implements X-AI to demystify a black-box CNN for processing surface EMG to conduct personal identification. The model’s attention is derived using Grad-CAM analysis, interpreting and extracting the unique neural code in a visualizable, and reportable manner.

### Contribution 2

This work specifically focuses on the dynamical transient phase of contraction, which includes imperative information regarding the users’ motor unit recruitment “patterns”. Different from the literature, which mainly includes the plateau phase of a contraction, our focus can provide more robustness while rejecting the magnitude sEMG artifacts.

### Contribution 3

The application of the proposed spatiotemporal mask generated based on the XAI outcomes reduces 40% of the input size and the model’s trainable parameters while compromising the performance by about 8%, pushing our research one step forward towards developing portable personal identification security hardware.

### Contribution 4

This paper explores the capability of sEMG signal space as a biomarker that uniquely and robustly identifies individuals on a large number and variety of gestures and subjects while including data from systematically selected sensors that detect activities across diverse muscle groups, maximizing identification accuracy.

### Contribution 5

This paper, for the first time, conducts a holistic optimization to find the minimal number of gestures and sensors. The gesture and sensor optimization not only maintains high performance of an identification system but also achieves high practicality that reduces the burden of tedious sEMG data collections and calibrations.

### Contribution 6

For model evaluation, instead of commonly used majority voting, which can ignore hidden false negative results, we use excessively stringent metrics to evaluate the model performance considering the predictions on every single short sliding window of spectrogram.

The rest of the paper is written as follows. Section II introduces the data acquisition and preprocessing. Section III provides details on the methods, including initial model architecture, gesture and sensor selection, and Grad-CAM analysis. The results are presented in Section IV. Section V highlights the superiority of our proposed model over commonly-used classic and DL models. Lastly, concluding remarks are provided in Section VI.

## II. Biometric Database

### A. Data Acquisition Process

Ninapro DB2 is used in this project. Ninapro is a publicly available open-source database utilized in this paper to evaluate the efficacy of the proposed methodology. For collecting the data set, Delsys Trigno system with 12 wireless electrodes is used, out of which eight channels are placed around the forearm near the radio-humeral joint, two are placed near the wrist on the extensor digitorum and flexor digitorum superficialis muscles, and two are placed on the biceps and triceps brachii muscles [25]. Fig. 1 shows the electrode placements.

**Fig. 1:**
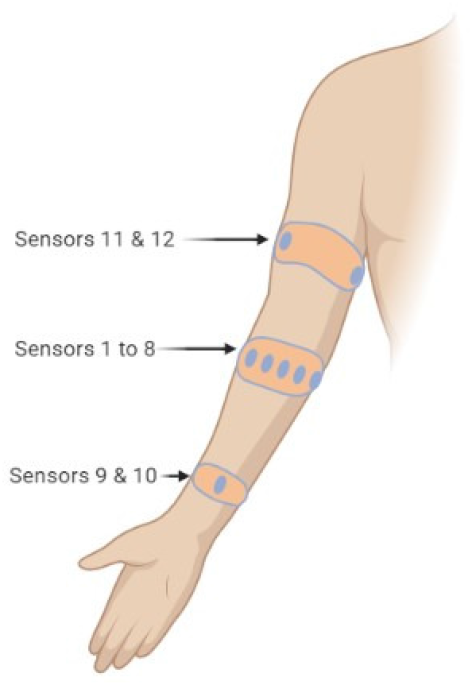
Placement of myoelectric sensors.

The sEMG signals are recorded from 40 intact subjects (12 females and 28 males) having age 29.9 ± 3.9 years. DB2 is segmented into three exercises: Exercises B, C, and D. Exercise B contains 17 gestures, among which 8 are various isometric and isotonic hand configurations, and 9 are wrist movements. Exercise C contains 23 gestures of grasping everyday objects and other functional movements. Exercise D contains 9 force patterns. We combine Exercises B (17 gestures) and C (23 gestures) for our research, taking into account a total of 40 gestures from 40 subjects. Subjects performed each hand movement 6 times, holding the gesture for 5 seconds followed by a rest of 3 seconds. Surface sEMG signals are sampled at a frequency of 2000 Hz. The motion labels are further refined [25].

### B. Data Preprocessing and Normalization

In this paper, we use *μ*-law transformation [26], [27], which is a logarithmic and nonlinear transformation. *μ*-law transformation is followed by z-score normalization, forming the proposed signal preprocessing pipeline. For z-score normalization, the mean and standard deviation are found from training data. Specifically let *i* be the sensor index, and 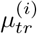 and 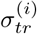 be the mean and standard deviation of the sEMG signals from sensor *i* of only the training data. The z-score normalization is defined as

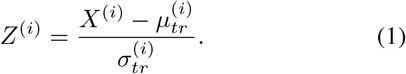

The *μ*-law transformation increases the distinguishability among sensors and has been widely used in speech processing. The *μ*-law transformation follows the mathematical design given by

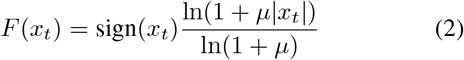

where *x_t_* denotes a single data value and *μ* = 2048 is used in this paper.

The duration of hand movement for different gestures and different subjects varies significantly over repetitions. These variations are often not considered in the literature; however, ignoring these variations results in bias in the data set. Thus, we only keep the first 1.5 seconds of data for each repetition that contains the transient and static part of the hand movement. *Taking into account the transient part of contractions allows us to potentially detect differences in dynamic motor unit recruitment during contraction, which can be a distinguishing factor for the understudied problem.* As a result, all subjects are represented with the same length of transient signals in the data set. The sEMG signal for each channel is segmented into windows, each having a length of 600 ms and a stride of 100 ms. The windowing process is demonstrated in Fig. 2. The 1D sEMG signal from each electrode is converted into 2D spectrogram images using Short Time Fourier Transform (STFT), which is applied with a window size of 500 ms and an overlap of 95%. The raw spectrogram (shown in Fig. 3a) is clipped at 500 Hz (as can be seen in Fig. 3b). Thus, the final shape of the 2D window (for each channel) after applying STFT and clipping can be represented by a 250 × 25 spectrogram, which is used later in this paper as the input to the CNN model. The processing pipeline is summarized in Fig. 4.

**Fig. 2:**
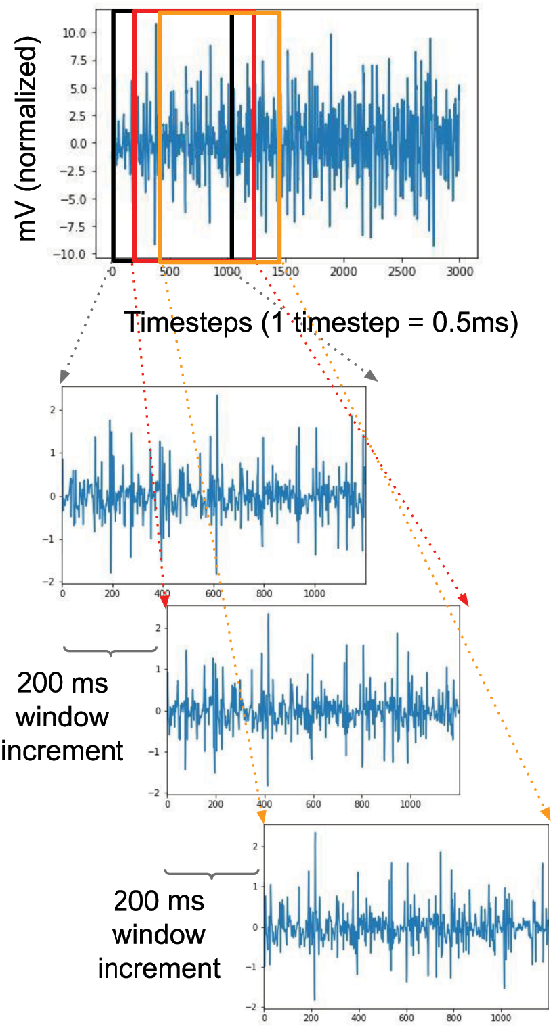
Segmentation of sEMG signal into small windows.

**Fig. 3:**
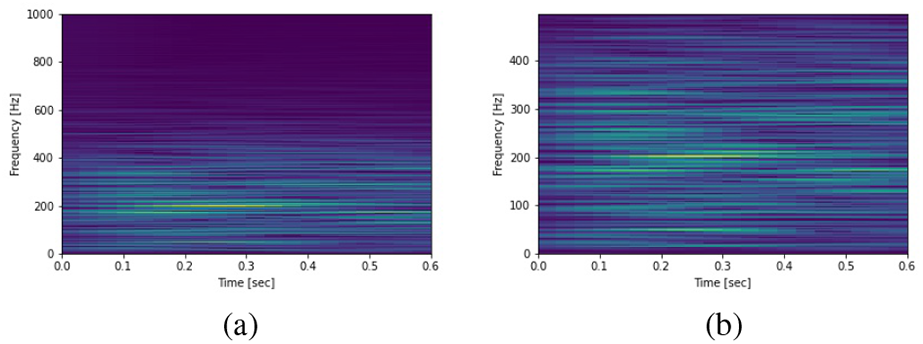
(a) Spectrogram of a channel, (b) Spectrogram after clipping high frequencies.

**Fig. 4:**
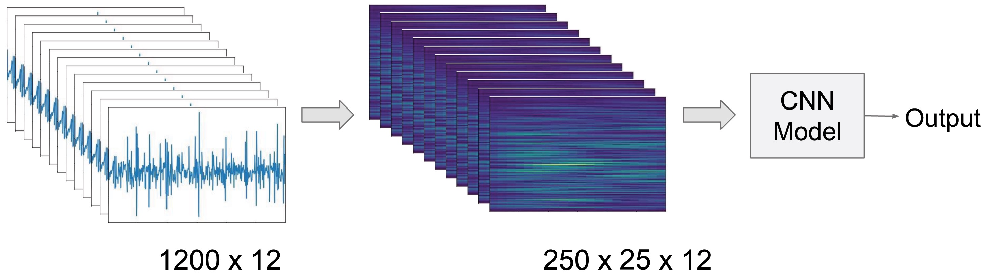
High-level overview of the complete process.

## III. Method

This paper aims to design a compact, robust, and explainable identification system. An identification system extracts user-specific transient feature patterns to detect one user among a database of pre-collected templates of many users. Ninapro DB2 allows us to explore the possibility of sEMG as a biomarker without the restriction of extracting neurophysiological patterns from a limited number and types of gestures and subjects. However, there is a trade-off between the system’s practicality and performance. Hence, finding the minimal gestures and sensors is critical to lift the burden of tedious data collection from users before the system is deployed. Applying representative feature-based clustering and individual performance ranking, we propose to optimize the number of gestures and sensors for training. Furthermore, this paper implements XAI to decode the model’s attention to investigate the spectrotemporal features (e.g., frequency ranges) that qualify sEMG as a biomarker. The details of the methodologies can be found in Subsections III-B and III-C.

### A. Initial Model Architecture

As discussed in Section II-B, a short window of sEMG signals is converted to a 2D image using STFT. The resulting images (the third dimension corresponds to the sensors) should be processed using the proposed neural network to detect the corresponding subject through a classification scheme. For this purpose, we exploit the power of neural networks for classification. Our model consists of two modules: the autonomous feature extractor and the classifier. The feature extractor should be able to capture the underlying features in the 2D representation of the sEMG signal. Therefore, CNNs are chosen as they have been shown to be powerful for feature extraction [28], [29]. Performing weight sharing through sliding kernels in a CNN results in a smaller number of trainable parameters. Also, sliding a kernel in 2D space, the CNN can detect the feature patterns appearing anywhere in a spectrogram. This lightweight but robust model structure is well-suited to our design of a compact identification system.

In this paper, we first tune the hyperparameters of the system and then discuss the model generalization and robustness (See Appendix A for more details). The proposed model contains four CNN blocks, each having a CNN layer (feature extractor) and a Rectified Linear Unit (ReLU) activation function, followed by a fully connected (FC) classifier (Fig. 5). In order to improve the model convergence during training, a 2D batch normalization is used in the last CNN block. The summary of the model is reported in Table I.

**Fig. 5:**
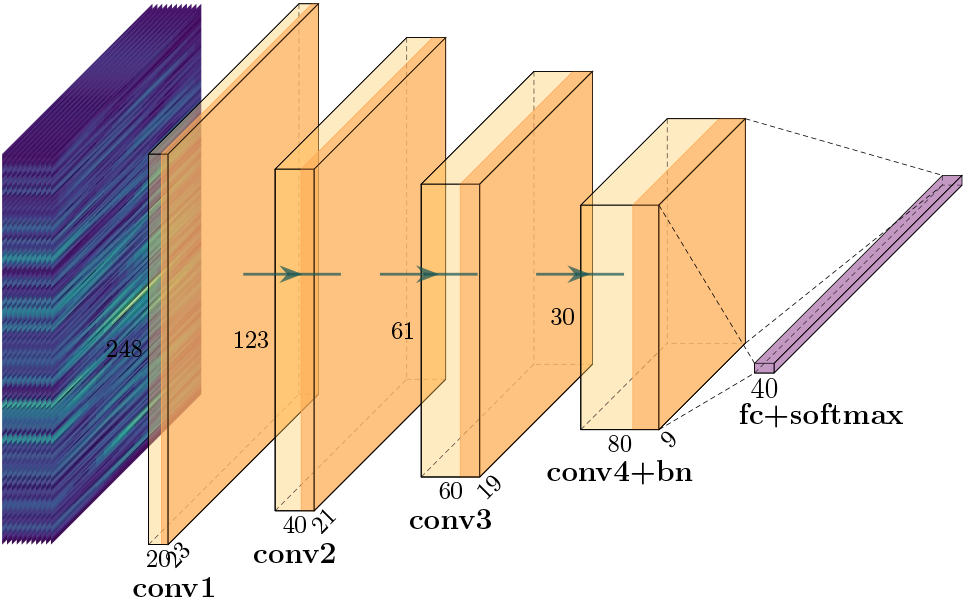
Overall architecture of the model.

**TABLE I:** Model architecture.

| Layer Name | Layer Type | # of Channels | Filter Size | Stride | Activation |
| --- | --- | --- | --- | --- | --- |
| conv1 | Conv2d | 20 | $3 \times 3$ | $1 \times 1$ | ReLU |
| conv2 | Conv2d | 40 | $3 \times 3$ | $2 \times 1$ | ReLU |
| conv3 | Conv2d | 60 | $3 \times 3$ | $2 \times 1$ | ReLU |
| conv4 | Conv2d | 80 | $3 \times 3$ | $2 \times 2$ | ReLU |
| bn | BatchNorm2d | 80 | - | - | ReLU |
| fc | FC | 21600 | - | - | - |

### B. Gesture and Sensor Selection

In order to enhance the practicality and usability of the proposed approach, we investigate how to reduce the number of (a) training gestures and (b) sensors to find the optimal selection while yielding similar performance to the larger number of gestures and sensors. The gesture-based and sensor-based optimization of input spaces are explained below.

The intuition of gesture-based optimization of the input space comes from synergistic similarities among sEMG signals of different gestures that can be clustered into lowdimensional finite groups [30], from each of which one representative gesture can be selected as a training gesture. The sensor-based optimization of the input space is based on individual sensor performance ranking. It should be noted that we conducted gesture and sensor selection in a sequential manner, meaning that for the gesture-based optimization, sEMG signals from all sensors are considered, while for the sensor-based optimization, the best gesture from the previous step is considered.

#### 1) Gesture-based Input Space Optimization

For gesturebased optimization, we extract various features from both time and frequency domains to capture distinguishing spectrotemporal patterns, which can be potentially different among selected gestures. The temporal features are Mean Absolute Value, Variance, Mean Square Root, Root Mean Square, Log Detector, Waveform Length, Difference Absolute Standard Deviation Value, Zero Crossing, Skewness, and Kurtosis [23]. For the spectral features, we considered conventional neural frequency bands of Delta (0.5-4Hz), Theta (4-8 Hz), Alpha (8-12 Hz), Beta (12-35 Hz), and Gamma (>35 Hz) [31], [32]. For each mentioned frequency band, mean power density is considered to be the spectral feature. We extract these features from each sensor on the sEMG signals after *μ*-law transformation and averaged over subjects and repetitions. Thus, the pre-clustering data dimensions in the time domain are 40 × 120 (where 40 corresponds to the number of gestures and 120 corresponds to the ten temporal features calculated for every 12 sensors). Also, in the frequency domain, the dimension is 40 × 60, where 40 represents the number of gestures and 60 corresponds to the five spectral features calculated for all 12 sensors.

GMM [33], [34] is used to conduct clustering for the gestures based on the extracted temporal and spectral features (180 for gesture clustering). In this paper, GMM is initialized using K-Means to increase the convergence speed and relieve some computational intensiveness. Compared to K-means, which gives each data point (e.g., a gesture) a hard assignment to a particular group, GMM is a soft clustering approach that gives a probability to each data point belonging to a Gaussian component. Furthermore, the GMM parameters (mixture weights, means, and variances) are iteratively updated through the expectation-maximization (EM) algorithm to find the maximum likelihood of a GMM best capturing the distribution of the gesture representations. A GMM represents the distribution of gestures in the form of

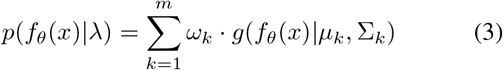

where *m* is the number of the user-defined clusters, *f_θ_*(*x*) is the representation of the gesture x, and *θ* is the parameter of the representation. λ is the vector of GMM parameters including mixture weights (*ω_k_*), mean vectors (*μ_k_*), and covariance matrices (∑_*k*_). Gaussian densities (*g*(*f_θ_*(*x*)|*μ_k_*, ∑_*k*_)) are given by

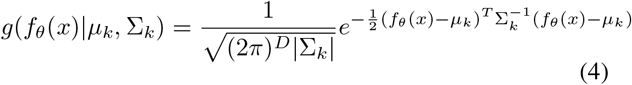

where *D* is the dimension of the representation.

Before implementing GMM, we first apply Principal Component Analysis (PCA) [35] to reduce the 40 × 180 feature space (containing 40 gestures with temporal and spectral features) to 40 × 15 for better clustering results. GMM clustering requires the number of clusters *m* as input. We implement widely adopted Bayesian Information Criterion (BIC) (see (5)) to derive the optimal number of clusters as the minimal number of training gestures [36].

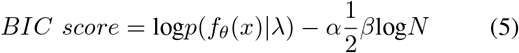

*α* is a penalty weight, *β* is the number of parameters in a GMM model, and *N* denote the number of gestures.

A GMM becomes more complex when the number of Gaussian components increases, potentially resulting in an overfitting problem. As a BIC score is penalized by the model complexity (the number of components) in a GMM, we choose the number of clusters as seven that has the lowest BIC score to avoid overfitting problems (see Fig. 6). The GMM result shows that seven Gaussian components can optimally and sufficiently capture the distribution of the 40 × 15 feature space. This paper selects one gesture with the highest log-likelihood from each assigned cluster as the representative training gesture of that cluster. As a result, we select seven optimal gestures (shown in Fig. 7a) to reduce the input space.

**Fig. 6:**
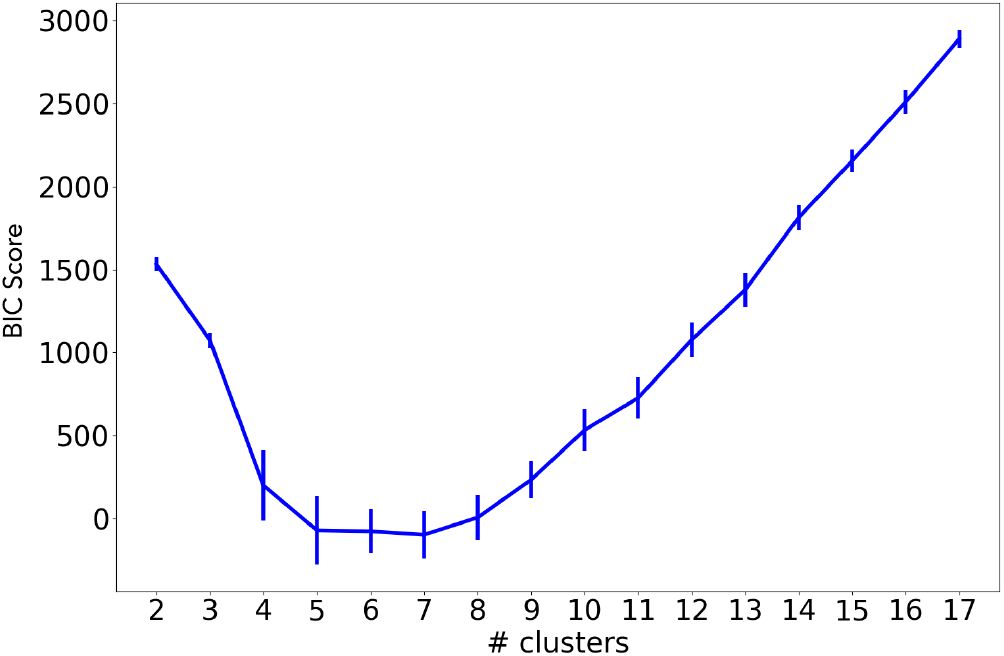
BIC scores for the derivation of the optimal number of clusters.

**Fig. 7:**
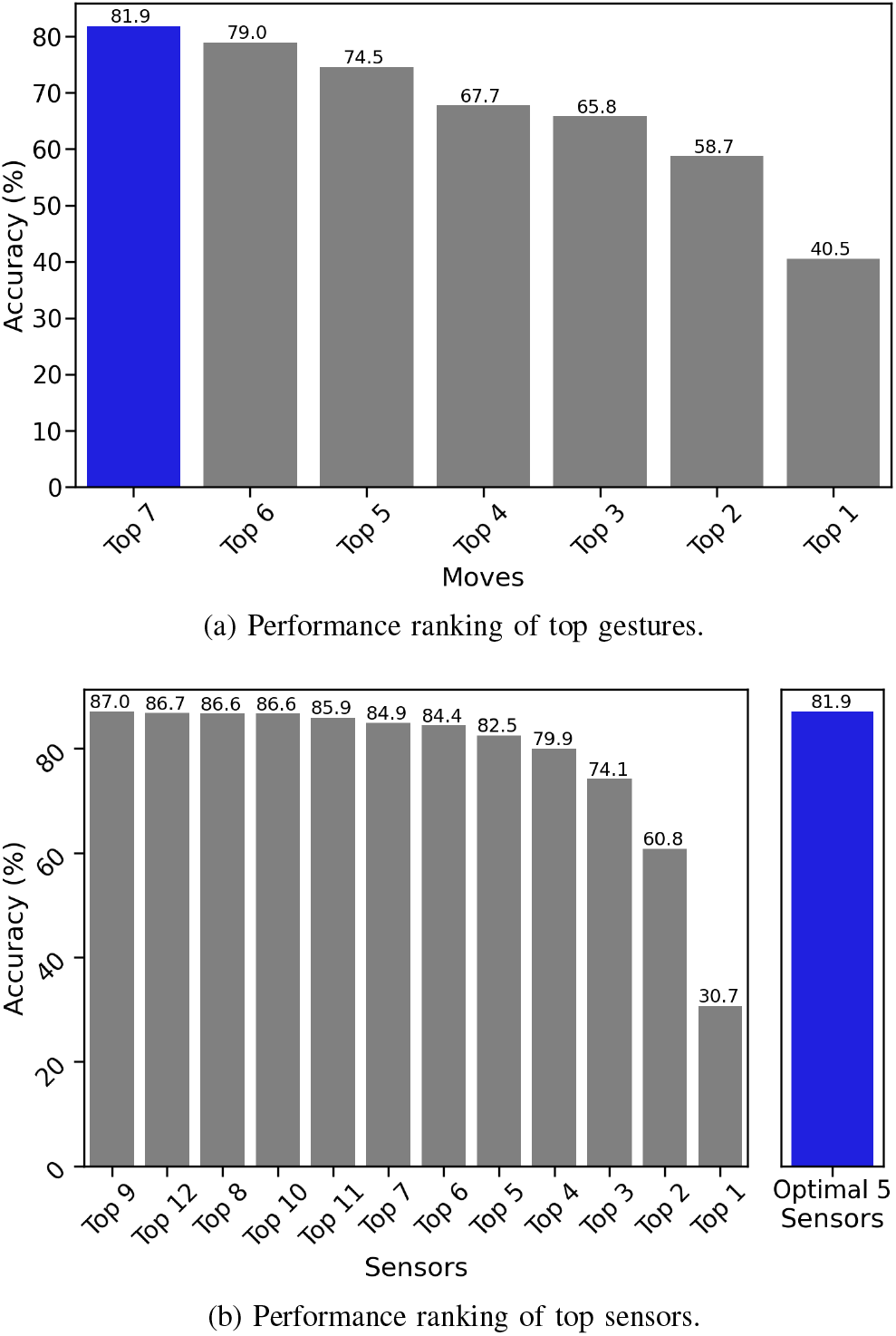
(a) Performance ranking of top gestures is based on the selected five optimal sensors; the blue bar shows the accuracy of the selected seven optimal gestures. (b) Performance ranking of top sensors is based on the selected seven optimal gestures; the blue bar shows the accuracy of the selected five optimal sensors.

#### 2) Sensor-based Input Space Optimization

The sensor optimization employs performance ranking to find the optimal sensors. In this approach, we feed sEMG signals from only one sensor at a time into the proposed model. The best-performing sensors are added one by one into the training set according to the individual performance in descending order for comparison. According to the One Standard Error Rule, performance ranking returns five as the optimal number of sensors for training, securing almost similar performance with smaller input size.

### C. Grad-CAM Analysis: Explainability-based Optimization

Grad-CAM has enhanced the transparency and explainability of a black-box CNN-based network through the gradients of any given class flowing into the last convolutional layer of the network, producing a heatmap that highlights the network attention on the input [37]. As a part of XAI, Grad-CAM is often used to reveal the attention of machine intelligence, to extract the underlying information that is invisible to the naked eyes, and to optimize the size of the data set and model architecture. In this paper, we utilize Grad-CAM to (1) help visualize the attention of the network on the average subject-wise spectrograms and the corresponding localization maps in parallel, (2) extract the identification code from the overlay of the averaged spectrogram and attention heatmap for each user, and (3) extensively reduce the input spaces and the number of the trainable model parameters, optimizing the size of the proposed network.

#### 1) Subject-wise Attention Heatmap Generation

In this paper, we concatenate the best five sensors horizontally to preserve the critical channel-wise localization information and show the model’s attention on different sensors. The Grad-CAM analysis is conducted on the best-performing model trained on the optimal gestures (seven gestures) and concatenated best sensors (five sensors) in the inference phase to demystify the model decision. The horizontal concatenation broadens the input size of each channel (the third axis of inputs) from 250 × 25 to 250 × 165 with a zero padding of 250 × 10 in between sensors to generate distance between each sensor information. We do not want the proposed model to treat transitions as part of the signal patterns; hence, we have introduced gaps using zero padding. Furthermore, the sensor concatenation and zero padding broaden the input size. To achieve better model performance, we slightly modify the proposed model architecture by setting the stride as 2 × 2 in all convolutional layers. The gradients from the last convolutional layer are extracted and resized from 30 × 19 to align with the input size to form the attention heatmap. Each heatmap indicates the model attention on each sample spectrogram. The average spectrograms and corresponding heatmaps by subjects are presented in parallel as the results of the attention analysis.

#### 2) Identification Code Extraction by Subject

By the visual analysis of Grad-CAM, the model attention varies in sensors and frequency ranges for different subjects, later defined as frequency bins. It indicates the distinguishability and uniqueness of the spectrotemporal neurophysiological characteristics of each subject. These underlying sEMG features can be translated into the identification codes that can also be put into practical use, such as personal authentication hardware.

To extract the unique identification code for each individual, we segment the average spectrogram of each subject ranging from 0 Hz to 500 Hz into 25 frequency bins, each containing 20 Hz spectrotemporal information across sensors. Thus, an identification code is an array of five scalar values, named as identification scalar, each falling into a range between 1 and 25. An identification scalar is calculated through the equation given by

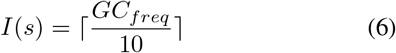

where *GC_freq_* is the coordinate of a gravity center on the frequency axis of a heatmap and *s* = 1,…, 5 is the index of the five optimal sensors. The function *center_of_mass* [38] is used to obtain the gravity center coordinates of the subject-wise Grad-CAM heatmaps of the sensors for each subject. We use the blurring and thresholding method to ensure the true gravity center at each sensor is precisely defined, reducing the noise. As the next step, we convert the heatmap to binary images to accentuate the hottest areas. It should be noted that this approach also avoids gravity center shifts. In Fig. 8, before applying blurring and thresholding, the gravity center of the given sensor is found between 151 and 160 on the frequency axis, translated into Code 16. However, after applying blurring and thresholding, the gravity center of the same sensor is found between 171 and 180 on the same frequency axis, translated into Code 18. Thus, we observe a shift of two in Code when the blurring and thresholding method is not applied.

**Fig. 8:**
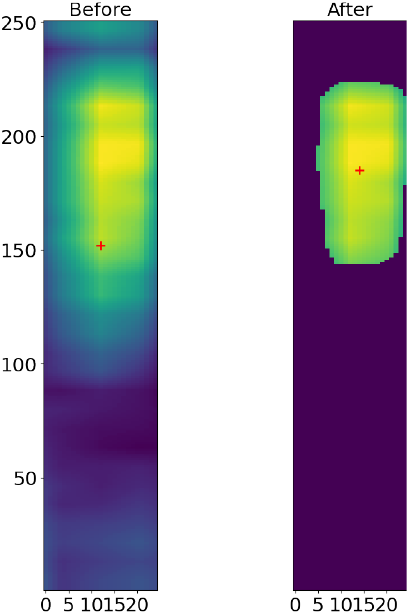
Heatmap of a sensor before and after blurring and thresholding, Subject 30 on Sensor 8.

However, this approach may fail to detect any hot zone when the heatmap is highly dispersed. To address the issue, the sensor-wise segmentation of the attention heatmap is used as an alternative approach that further divides the heatmap corresponding to the sensor into five equal 100 Hz segments. Each segment returns an average heat, indicating the strength of the model’s attention on that segment. By using the same function, five gravity center candidates can be found at each sensor. The gravity center of the segment that has the highest average heat is considered as the final candidate for the gravity center at the sensor. Thus, the gravity centers respectively represent the most concentrated attention spots for the best sensors on each heatmap. The resulting five centers form the identification code of a particular subject, which shows the specific attention of the network on different frequencies and sensors for identifying each subject.

The radius of an identification code calculates the Euclidean norm of that identification code through

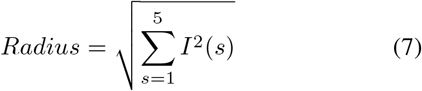

*I*(*s*) is an identification scalar.

This measurement indicates the main frequencies where the proposed model pays the most attention when predicting a subject. The identification code radius of each subject is normalized to a range between 1 and 25 to be consistent with the frequency bin numbers. The radius distribution analysis evaluates the relation between the model attention (on frequency bins) and the model performance (on each subject group in descending order), as shown in Fig. 9. The analysis results show that the proposed model pays attention to wide-ranging frequency bins, especially including higher frequencies (higher gamma band of >80 Hz) for best-performing subjects, serving as a reference for the mask generation in the next subsection.

**Fig. 9:**
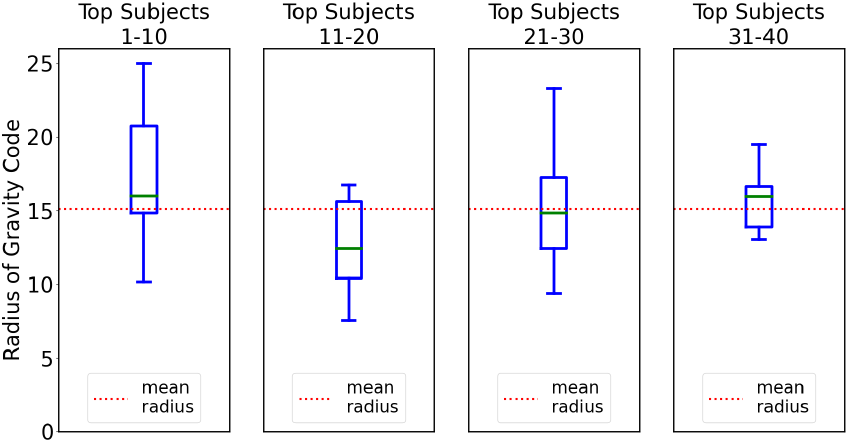
Radius distribution of identification code by top-performing subjects. Header “Top Subjects 1-10” means the Top-1-to-10-performing subjects. The meaning of the other three headers follows the same pattern.

#### 3) Attention-based Spectrotemporal Mask Generation and Model’s Size Optimization

Based on the previously mentioned gesture-based and sensor-based optimizations, we enhance the practicality by minimizing the number of gestures and sensors used in training. The smaller input space and less trainable parameters can further refine the proposed identification system by reducing the data storage, speeding up the training process, and increasing the practicality.

In this subsection, we propose an optimizing attention-based spectrotemporal mask that abandons the trivial areas which play the minimum role in classification from the input space. We hypothesize that re-training the model only on the most informative segments of spectrograms can result in similar performance while significantly reducing the model size. The median Grad-CAM heatmap of the top-10-performing samples of the spectrograms from the test set is utilized to generate the most significant attention-based segment of spectrotemporal information across subjects and gestures. The median heatmap rather than the average heatmap is employed for the mask generation as the data may not be normally distributed. Based on the results achieved on the model attention summarized from the previous radius distribution analysis, the optimizing spectrotemporal mask is systematically calculated using sensor segmentation. We segment each sensor into multiple fine pieces and select the top 60% segments that have the highest average heat. The outcome consists of both low- and high-frequency areas at each sensor (Fig. 10). It is expected that our model pays attention to both low- and high-frequency areas on the average spectrogram because our input signals include low-frequency contraction at transient phase and high-frequency contraction at plateau phase.

**Fig. 10:**
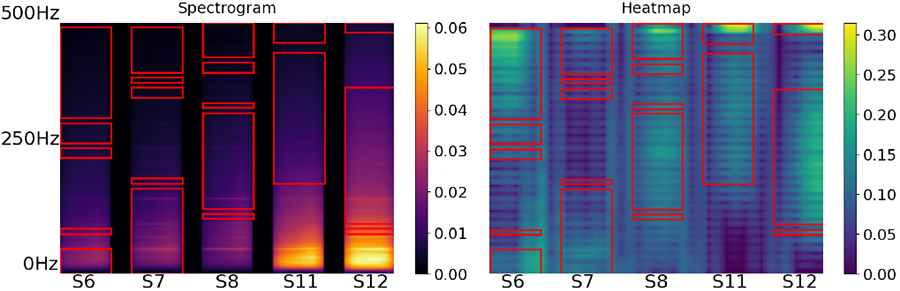
Spectrogram segmentation and mask generation.

After applying the mask (calculated based on the average attention map of best-performing subjects) on each spectrogram for all subjects, the segments for each sensor are concatenated vertically, making one transformed spectrogram for each sensor. The resulting five transformed spectrograms are horizontally concatenated with zero paddings in between, forming the small input space (see Fig. 11) for the model. The model is re-trained on the reduced data set.

**Fig. 11:**
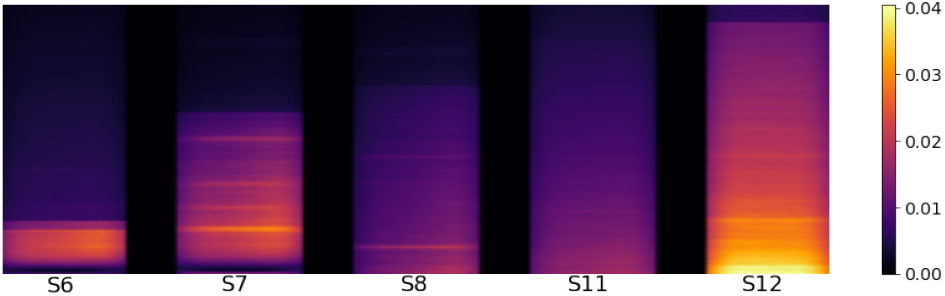
Example: mask application result.

## IV. Results

In all the experiments, the models are trained for a maximum of 500 epochs with a batch size of 32. Adam optimizer is used with a learning rate of 0.0001, which is reduced by a factor of 0.1 after the first 100 epochs.

### A. Gesture and Sensor Selection

In gesture-based optimization, we investigate five, six, and seven gestures and evaluate the corresponding model performance given all 12 sensors because of the similar BIC scores. According to the One Standard Error Rule, we choose the seven optimal gestures of 4, 12, 15, 22, 26, 30, and 32 based on the mixed-domain (temporal and spectral) clustering, achieving an accuracy of 86.735%.

In sensor-based optimization, we derive optimal sensors based on the ranking of individual sensor performance and then combine the most informative sensors. An identification system is easier to use when a user is required to attach sensors to fewer locations on the arm, potentially attracting more users. In order to further enhance the practicality of our identification system, our analysis results in Sensor IDs, 6, 7, 8, 11, and 12 to be the best five sensors spreading among two muscle groups (extensor-flexor group and biceps and triceps group), achieving the same accuracy as the top 5 sensors (see Fig. 7b). By running the model on these five sensors (which contain the maximum information when compared with the case of having 12 sensors for the chosen gestures), the accuracy of 81.866% was observed, which shows only less than 5% compromise. **It should be highlighted that the gesture and sensor optimization reduce the input size of the model by 85%**, compared to the input size when conducting hyperparameter tuning (see Table. III).

### B. Grad-CAM

Fig. 12 shows the spectrograms and attention heatmaps (generated by GRAD-CAM) of two Subjects (i.e., #19 and #28) with top performance. We can conclude that the proposed model makes decisions based on distinct frequency bins among sensors which reveals the underlying spectrotemporal patterns that can be exploited to identify subjects and optimize the proposed model. As explained, the unique features of the attention heatmaps are translated into identification codes, each consisting of five frequency bin numbers ranging from 1 to 25. For example, the algorithm generates the identification codes of 11-15-23-4-1 for Subject #19, and 25-25-25-12-6 for Subject #28.

**Fig. 12:**
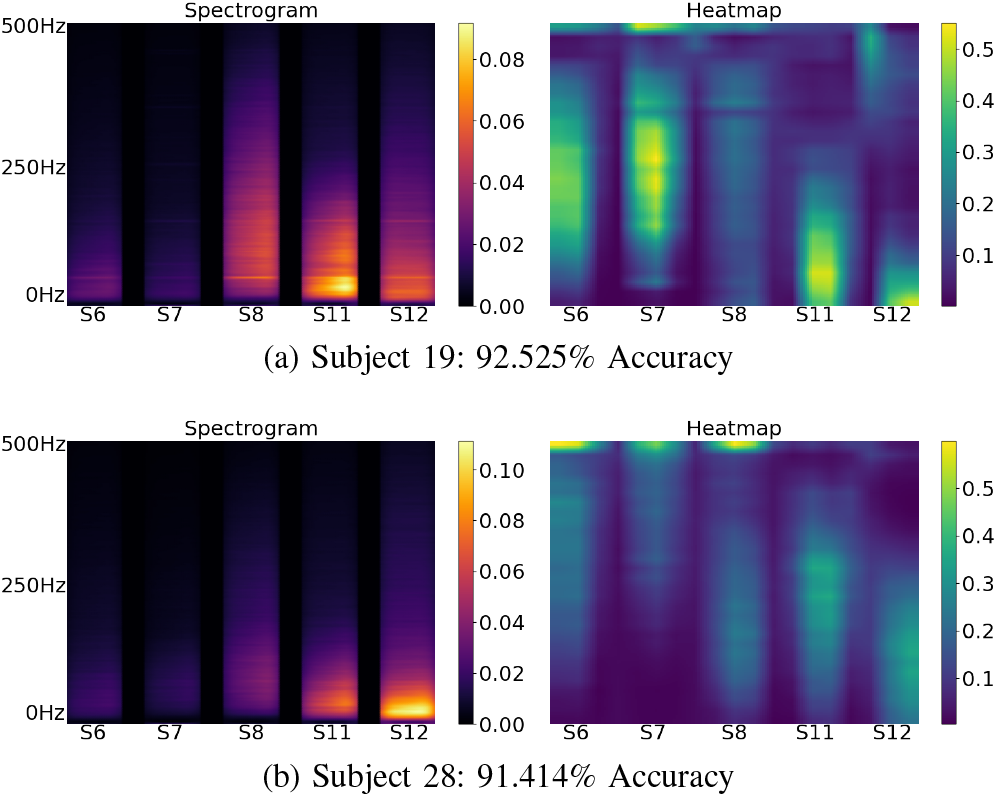
Average spectrogram and heatmap of top-performed subjects #19 and #28. The left figure is the spectrogram, and the right figure is the attention heatmap.

As mentioned, we utilize GradCAM to further reduce the size of the network by optimizing the input space. The results show that the application of the spectrotemporal mask discards 40% of the individual input size from 250 × 165 to 150 × 165. Hence, the proposed approach allows for dropping 40% of the trainable parameters of the network, from 938K to 563K parameters, resulting in a much less complex network. Also, it results in over 20% reduction of time needed for training (this number may vary on different machines). The approach achieves all of these while compromising about 8% of accuracy compared with the model performance in the optimization section (see Subsection IV-A). This shows the efficacy of the proposed attention-based data masking optimization technique proposed in this paper.

### V. Comparative Study

The goal of this comparative study is to highlight the superiority of our proposed CNN model over the commonly used classic and deep learning models when training on the optimal gestures and sensor derived in Section III-B. Thus we compare our proposed model with (a) a two-layer multilayer perceptron (MLP) model, (b) a two-module hybrid model with four CNN blocks followed by six LSTM layers and a fully connected layer, and (c) and a classic SVM model. In this comparative study, each comparing model trains on the seven optimal gestures (4, 12, 15, 22, 26, 30, and 32) and the most informative five sensors (6, 7, 8, 11, and 12). The validation data includes sEMG signals from the even repetitions (2, 4, and 6) of the remaining 33 gestures, while the test data contains the sEMG signals from the odd repetitions (1, 3, and 5) of the same 33 gestures.

In this Section, we select the comparing models (neural networks) to be structurally comparable to our proposed CNN model. The MLP model has 30 neurons on the hidden layer. We modified our recently proposed hybrid model [39] specifically for the identification problem. The hybrid model has a CNN module followed by an LSTM module. The CNN module consists of four CNN blocks, each having a 2D convolutional layer, a Batch Normalization layer, and a ReLU layer. The convolutional layers have 20, 40, 60, and 80 channels, respectively, with a kernel size of 3 × 3. The output from the last CNN block is fed to the LSTM module, which consists of six LSTM layers with 92 hidden units on each layer. The last layer is a fully connected layer with 4240 neurons. For the SVM model, we extracted Mean, Median, Root Mean Square, and Variance from sliding windows of size 25 × 10 with a 20% overlap along the frequency axis on each input spectrogram. This results in a feature vector of size 80 for each sensor. This procedure is done for all 12 sensors. Therefore, each sample spectrogram is converted to a vector of 960 features. The results are summarized in Table II and highlighted in the following contributions.

**TABLE II:**
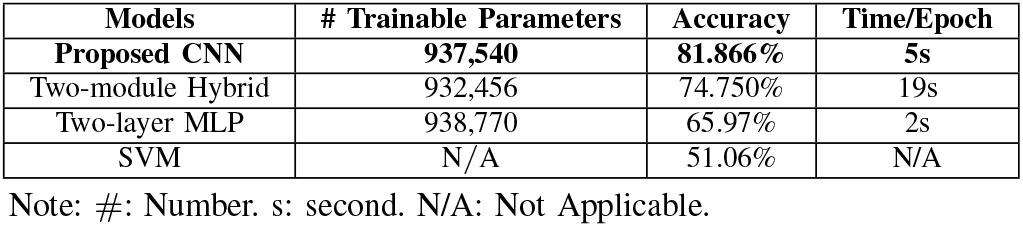
Results for comparing the proposed model with commonly used classic and deep learning models.

| Models | # Trainable Parameters | Accuracy | Time/Epoch |
| --- | --- | --- | --- |
| **Proposed CNN** | **937,540** | **81.866%** | **5s** |
| Two-module Hybrid | 932,456 | 74.750% | 19s |
| Two-layer MLP | 938,770 | 65.97% | 2s |
| SVM | N/A | 51.06% | N/A |

Note: #: Number. s: second. N/A: Not Applicable.

#### Observation 1

The SVM fails to secure accuracy of above 70%. The hybrid model achieves about 75% accuracy, which is 7% lower than our proposed CNN.

#### Observation 2

The MLP also fails to secure above 70% accuracy. The comparing MLP model architecture has to be simple (two layers) to match the structural complexity of our proposed model. Rather than flattening the inputs, training a CNN model preserves the spatiotemporal information of the spectrograms. Leveraging kernel sliding, our CNN model can detect neural feature patterns appearing anywhere in a spectrogram based on a smaller amount of training data than the data needed for training an MLP for the same task.

#### Observation 3

Our CNN model is trained 74% (or 5/19) faster than the hybrid model on each iteration. Given the similar model convergence, which is the number of training iterations for a model to achieve its maximum performance, our proposed model is more efficient than the hybrid model.

Considering the above observations, the proposed model proves to be considerably well-suited with a compact, practical, explainable, and robust design for personal identification systems.

### VI. Conclusion

In this paper, we investigate the possibility of using the hidden underlying neurophysiological patterns in multichannel surface electromyography signals to identify users while securing a high performance. We propose and evaluate an optimized and explainable neural network that analyzes the information context of gestures and sensors to find out the minimal but sufficient number of best gestures, best sensors, and best frequency bands for training the model to enhance practicality and efficiency. We have shown that the performance can be preserved using data from only two muscle groups. The Grad-CAM analysis is also performed to decode the attention of the neural network model. The outcome of Grad-CAM analysis is also utilized to significantly reduce the needed data size and thereby reduce the number of trainable parameters of the model, reducing the complexity and increasing the speed of training. This paper sheds light on the capacity of the underlying neurophysiological signature of sEMG biosignals for identifying individuals. XAI helps visualize the unique and complex neural feature patterns associated with each subject and quantify these patterns through identification code, pushing forward the biometric research on human identification. In order to further enhance the model’s generalizability and robustness, collecting sEMG signals from more subjects performing more gestures in multiple sessions could be one of our future lines of research.

## Appendix A Model Optimization

During our experiments, we have divided the data set into three subsets called training, validation, and test set. The validation set is used to pick the best model during training on the training set. The hyperparameter tuning is done in two steps. First, we feed as much as information available to the model and validating its performance (named as Problem 1). For this purpose, we have considered all the gestures for the three sets of data, but they differ in repetitions. In other words, the subjects performed a set of gestures during training and repeated the same gestures during the inference phase. In the second step of tuning, we have considered a more challenging problem in which the set of gestures used to train the model is entirely isolated from the one used to validate and test the model during the inference (named as Problem 2). Therefore, we picked 18 gestures for the training set, and the remaining gestures (e.g., 22) are used for validation and test sets. Since the gestures set for the validation and test set are the same, different repetitions are considered for each of them. Moreover, to show the independence of our results to the choice of the training, validation, and test sets, we have considered various random settings for each step. For step one, two settings with randomized repetitions are considered (Table III). For the second step, gesture sets as well differ, which resulted in the four settings shown in Table IV. Corresponding accuracies are reported in Table V. It could be seen that the results are consistent.

**TABLE III:** Problem 1: train-validate-test settings.

|  | Train |  | Valid |  | Test |  |
| --- | --- | --- | --- | --- | --- | --- |
|  | Gestures | Rep. | Gestures | Rep. | Gestures | Rep. |
| Set 1 | All 40 | 2, 4, 6 | All 40 | 1 | All 40 | 3, 5 |
| Set 2 | All 40 | 1, 3, 6 | All 40 | 2 | All 40 | 4, 5 |

Note: Gestures are from 1 to 40. Rep. denotes repetitions.

**TABLE IV:**
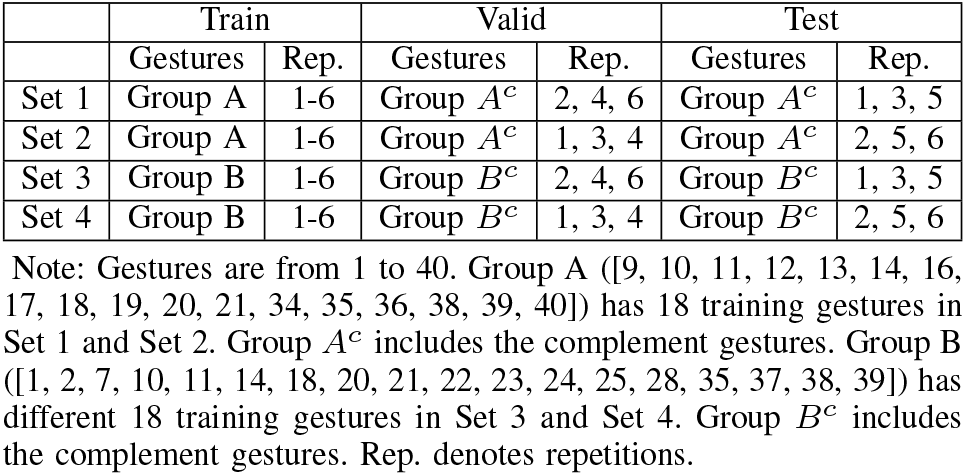
Problem 2: train-validate-test settings.

**TABLE V:** Model performance on train, validation, and test sets for each problem settings.

|  | Setting | Train | Validation | Test |
| --- | --- | --- | --- | --- |
| Problem 1 | Setting 1 | 99.621 | 97.244 | 97.9842 |
|  | Setting 2 | 99.642 | 98.075 | 97.703 |
| Problem 2 | Setting 1 | 99.866 | 88.492 | 88.879 |
|  | Setting 2 | 99.824 | 88.788 | 88.966 |
|  | Setting 3 | 99.528 | 91.705 | 90.875 |
|  | Setting 4 | 99.972 | 91.686 | 90.693 |

## References

[1] T. Yaqoob, H. Abbas, and N. Shafqat, “Integrated security, safety, and privacy risk assessment framework for medical devices,” IEEE Journal of Biomedical and Health Informatics, vol. 24, no. 6, pp. 1752–1761, 2020.

[2] A. Abbas and S. U. Khan, “A review on the state-of-the-art privacy-preserving approaches in the e-health clouds,” IEEE Journal of Biomedical and Health Informatics, vol. 18, no. 4, pp. 1431–1441, 2014.

[3] A. Khedr and G. Gulak, “Securemed: Secure medical computation using gpu-accelerated homomorphic encryption scheme,” IEEE Journal of Biomedical and Health Informatics, vol. 22, no. 2, pp. 597–606, 2018.

[4] R. Sánchez-Guerrero, F. A. Mendoza, D. Díaz-Sánchez, P. A. Cabarcos, and A. M. López, “Collaborative ehealth meets security: Privacyenhancing patient profile management,” IEEE Journal of Biomedical and Health Informatics, vol. 21, no. 6, pp. 1741–1749, 2017.

[5] J. Long, No tech hacking: A guide to social engineering, dumpster diving, and shoulder surfing. Syngress, 2011.

[6] W. Yang, N. Li, O. Chowdhury, A. Xiong, and R. W. Proctor, “An empirical study of mnemonic sentence-based password generation strategies,” in Proceedings of the ACM SIGSAC conference on computer and communications security, Vienna, Austria, Oct 2016, pp. 1216–1229.

[7] M. Cardaioli, M. Conti, K. Balagani, and P. Gasti, “Your pin sounds good! augmentation of pin guessing strategies via audio leakage,” in Proceedings of the European Symposium on Research in Computer Security. Guildford, United Kingdom: Springer, Sept 2020, pp. 720–735.

[8] S. Mamonov and R. Benbunan-Fich, “The impact of information security threat awareness on privacy-protective behaviors,” Computers in Human Behavior, vol. 83, pp. 32–44, 2018.

[9] K. Erickson and P. N. Howard, “A case of mistaken identity? news accounts of hacker, consumer, and organizational responsibility for compromised digital records,” Journal of computer-mediated communication, vol. 12, no. 4, pp. 1229–1247, 2007.

[10] S. Alrwais, K. Yuan, E. Alowaisheq, X. Liao, A. Oprea, X. Wang, and Z. Li, “Catching predators at watering holes: finding and understanding strategically compromised websites,” in Proceedings of the 32nd Annual Conference on Computer Security Applications, Los Angeles, CA, Dec 2016, pp. 153–166.

[11] S. Hadiyoso, S. Aulia, and A. Rizal, “One-lead electrocardiogram for biometric authentication using time series analysis and support vector machine,” (IJACSA) International Journal of Advanced Computer Science and Applications, vol. 10, no. 2, 2019.

[12] Y. Zhang and M. Juhola, “On biometrics with eye movements,” IEEE Journal of Biomedical and Health Informatics, vol. 21, no. 5, pp. 1360–1366, 2017.

[13] P. Hu, H. Ning, T. Qiu, H. Song, Y. Wang, and X. Yao, “Security and privacy preservation scheme of face identification and resolution framework using fog computing in internet of things,” IEEE Internet of Things Journal, vol. 4, no. 5, pp. 1143–1155, 2017.

[14] M. P. Yankov, M. A. Olsen, M. B. Stegmann, S. S. Christensen, and S. Forchhammer, “Fingerprint entropy and identification capacity estimation based on pixel-level generative modelling,” IEEE Transactions on Information Forensics and Security, vol. 15, pp. 56–65, 2019.

[15] L. Lu, J. Mao, W. Wang, G. Ding, and Z. Zhang, “A study of personal recognition method based on emg signal,” IEEE Transactions on Biomedical Circuits and Systems, vol. 14, no. 4, 2020.

[16] T. Matsumoto, H. Matsumoto, K. Yamada, and S. Hoshino, “Impact of artificial” gummy” fingers on fingerprint systems,” in Proceedings of the Optical Security and Counterfeit Deterrence Techniques IV, vol. 4677. San Jose, CA: International Society for Optics and Photonics, Apr 2002, pp. 275–289.

[17] V. Ruiz-Albacete, P. Tome-Gonzalez, F. Alonso-Fernandez, J. Galbally, J. Fierrez, and J. Ortega-Garcia, “Direct attacks using fake images in iris verification,” in Proceedings of the European Workshop on Biometrics and Identity Management. Roskilde, Denmark: Springer, May 2008, pp. 181–190.

[18] C. De Luca, “Electromyography,” Encyclopedia of medical devices and instrumentation, 2006.

[19] S. A. Raurale, J. McAllister, and J. M. del Rincon, “Real-time embedded emg signal analysis for wrist-hand pose identification,” IEEE Transactions on Signal Processing, vol. 68, pp. 2713–2723, 2020.

[20] X. Jiang, K. Xu, X. Liu, C. Dai, D. Clifton, E. A. Clancy, M. Akay, and W. Chen, “Cancelable hd-semg-based biometrics for cross-application discrepant personal identification,” IEEE Journal of Biomedical and Health Informatics, pp. 1–1, 2020.

[21] S. Shin, J. Jung, and Y. T. Kim, “A study of an emg-based authentication algorithm using an artificial neural network,” IEEE SENSORS, pp. 1–3, 2017.

[22] H. Yamaba, T. Kurogi, K. Aburada, S.-I. Kubota, T. Katayama, M. Park, and N. Okazaki, “On applying support vector machines to a user authentication method using surface electromyogram signals,” Artificial Life and Robotics, vol. 23, no. 1, pp. 87–93, 2018.

[23] Q. Li, P. Dong, and J. Zheng, “Enhancing the security of pattern unlock with surface emg-based biometrics,” Applied Sciences, vol. 10, no. 2, p. 541, 2020.

[24] R. Shioji, S.-i. Ito, M. Ito, and M. Fukumi, “Personal authentication and hand motion recognition based on wrist emg analysis by a convolutional neural network,” in Proceedings of the IEEE International Conference on Internet of Things and Intelligence System (IOTAIS). IEEE, Nov 2018, pp. 184–188.

[25] M. Atzori, A. Gijsberts, C. Castellini, B. Caputo, A.-G. M. Hager, S. Elsig, G. Giatsidis, F. Bassetto, and H. Müller, “Electromyography data for non-invasive naturally-controlled robotic hand prostheses,” Scientific data, vol. 1, no. 1, pp. 1–13, 2014.

[26] E. Rahimian, S. Zabihi, S. F. Atashzar, A. Asif, and A. Mohammadi, “XceptionTime: Independent Time-Window xceptiontime architecture for hand gesture classification,” in Proceedings of the IEEE International Conference on Acoustics, Speech and Signal Processing (ICASSP), May 2020, pp. 1304–1308.

[27] E. Rahimian, S. Zabihi, A. Asif, S. F. Atashzar, and A. Mohammadi, “Few-Shot learning for decoding surface electromyography for hand gesture recognition,” in Proceedings of the IEEE International Conference on Acoustics, Speech and Signal Processing (ICASSP), Jun. 2021, pp. 1300–1304.

[28] M. D. Zeiler and R. Fergus, “Visualizing and understanding convolutional networks,” in Proceedings of the 13th European Conference on Computer Vision. Zurich, Switzerland: Springer, Sept 2014, pp. 818–833.

[29] C. Szegedy, S. Ioffe, V. Vanhoucke, and A. A. Alemi, “Inception-v4, inception-resnet and the impact of residual connections on learning,” in Proceedings of the 31th AAAI Conference on Artificial Intelligence, San Francisco, CA, Feb 2017.

[30] G. Jia, H. K. Lam, S. Ma, Z. Yang, Y. Xu, and B. Xiao, “Classification of electromyographic hand gesture signals using modified fuzzy c-means clustering and two-step machine learning approach,” IEEE Transactions on Neural Systems and Rehabilitation Engineering, vol. 28, no. 6, pp. 1428–1435, 2020.

[31] P. A. Abhang, B. W. Gawali, and S. C. Mehrotra, Introduction to EEG-and speech-based emotion recognition. Academic Press, 2016.

[32] K. P. Thomas and A. P. Vinod, “Toward eeg-based biometric systems: The great potential of brain-wave-based biometrics,” IEEE Systems, Man, and Cybernetics Magazine, vol. 3, no. 4, pp. 6–15, 2017.

[33] S. Wang, G. Azzari, and D. B. Lobell, “Crop type mapping without field-level labels: Random forest transfer and unsupervised clustering techniques,” Remote Sens. Environ., vol. 222, pp. 303–317, Mar. 2019.

[34] J. Wang and J. Jiang, “Unsupervised deep clustering via adaptive GMM modeling and optimization,” Neurocomputing, vol. 433, pp. 199–211, Apr. 2021.

[35] J. Chen, G. Wang, and G. B. Giannakis, “Nonlinear dimensionality reduction for discriminative analytics of multiple datasets,” IEEE Transactions on Signal Processing, vol. 67, no. 3, pp. 740–752, 2019.

[36] M. Nishida and T. Kawahara, “Unsupervised speaker indexing using speaker model selection based on bayesian information criterion,” in Proceedings of IEEE International Conference on Acoustics, Speech, and Signal Processing, vol. 1, Hong Kong, China, Apr 2003, pp. I–I.

[37] R. R. Selvaraju, M. Cogswell, A. Das, R. Vedantam, D. Parikh, and D. Batra, “Grad-cam: Visual explanations from deep networks via gradient-based localization,” in Proceedings of the IEEE International Conference on Computer Vision (ICCV), Oct 2017, pp. 618–626.

[38] P. Virtanen, R. Gommers, T. E. Oliphant, M. Haberland, T. Reddy, D. Cournapeau, E. Burovski, P. Peterson, W. Weckesser, J. Bright, S. J. van der Walt, M. Brett, J. Wilson, K. J. Millman, N. Mayorov, A. R. J. Nelson, E. Jones, R. Kern, E. Larson, C. J. Carey, I. Polat, Y. Feng, E. W. Moore, J. VanderPlas, D. Laxalde, J. Perktold, R. Cimrman, I. Henriksen, E. A. Quintero, C. R. Harris, A. M. Archibald, A. H. Ribeiro, F. Pedregosa, P. van Mulbregt, and SciPy 1.0 Contributors, “SciPy 1.0: Fundamental Algorithms for Scientific Computing in Python,” Nature Methods, vol. 17, pp. 261–272, 2020.

[39] P. Gulati, Q. Hu, and S. F. Atashzar, “Toward deep generalization of peripheral emg-based human-robot interfacing: A hybrid explainable solution for neurorobotic systems,” IEEE Robotics and Automation Letters, vol. 6, no. 2, pp. 2650–2657, 2021.

